# Avoiding Data Loss: Synthetic MRIs Generated from Diffusion Imaging Can Replace Corrupted Structural Acquisitions For Freesurfer-Seeded Tractography

**DOI:** 10.1101/2021.02.08.430215

**Authors:** Jeremy Beaumont, Giulio Gambarota, Marita Prior, Jurgen Fripp, Lee B. Reid

## Abstract

1

Magnetic Resonance Imaging (MRI) motion artefacts frequently complicate structural and diffusion MRI analyses. While diffusion imaging is easily ‘scrubbed’ of motion affected volumes, the same is not true for structural images. Structural images are critical to most diffusion-imaging pipelines thus their corruption can lead to disproportionate data loss. To enable diffusion-image processing when structural images have been corrupted, we propose a means by which synthetic structural images can be generated from diffusion MRI. This technique combines multi-tissue constrained spherical deconvolution, which is central to many existing diffusion analyses, with the Bloch equations which allow simulation of MRI intensities given scanner parameters and magnetic resonance (MR) tissue properties. We applied this technique to 32 scans, including those acquired on different scanners, with different protocols and with pathology present. The resulting synthetic T1w and T2w images were visually convincing and exhibited similar tissue contrast to acquired structural images. These were also of sufficient quality to drive a Freesurfer-based tractographic analysis. In this analysis, probabilistic tractography connecting the thalamus to the primary sensorimotor cortex was delineated with Freesurfer, using either real or synthetic structural images. Tractography for real and synthetic conditions was largely identical in terms of both voxels encountered (Dice 0.88 – 0.95) and mean fractional anisotropy (intrasubject absolute difference 0.00 – 0.02). We provide executables for the proposed technique in the hope that these may aid the community in analysing datasets where structural image corruption is common, such as studies of children or cognitively impaired persons.

**Highlights:**

- We propose a simple means of synthesizing T1w and T2w images from diffusion data
- The proposed method worked well for a variety of acquisitions
- Synthetic images showed tissue contrast akin to acquired images
- Synthetic images were high enough quality to be used for Freesurfer seeded diffusion tractography
- This method enables analysis of datasets where motion has corrupted acquired structural MRIs

## 2 Introduction

Diffusion magnetic resonance imaging (MRI) can quantify certain aspects of tissue microstructure and can, via tractography, delineate axonal pathways from which such microstructural measurements can be taken. These abilities make diffusion MRI a popular means of identifying and quantifying brain reorganisation and injury. Currently, such analyses almost always require identification of cortical or subcortical structures using an aligned structural MR image, such as a T1- or T2-weighted scan. Such reliance on structural images can introduce four difficulties into analysis pipelines. Firstly, cross-modality registration sometimes fails due to the meaningful difference between tissue contrasts combined with the typically low (2 – 2.5mm) spatial resolution of diffusion images. Secondly, even after correction diffusion images can display spatial distortions that are not found in structural images which can prevent perfect registration. Thirdly, reliance on structural MR images provides a greater risk that motion artefacts will prevent analysis. Specifically, diffusion MR sequences have a reasonable tolerance for motion as motion-affected volumes can be rejected from the 4D series and still leave sufficient information for analysis [1]. Structural images, however, are easily corrupted by such motion and cannot easily be ‘scrubbed’ in the same way. This issue is particularly prevalent in young children and populations with brain injury, who are highly valuable participants but also highly likely to move during scanning [2] and often less tolerant to remaining in the scanner for repeat scans. Finally, diffusion acquisitions are typically limited to lower spatial or angular resolution than desired due to scan time requirements. The requirement for high-quality structural images reduces the time available for diffusion acquisitions and thus their quality. This particularly affects studies of populations where motion artefacts are commonplace because, for such studies, additional scan time must be pre-allocated for the re-acquisition of motion affected structural images.

In this study we demonstrate a straightforward means of generating synthetic T1w and T2w images from a typical diffusion acquisition. This process takes advantage of recent advances that allow approximate calculation of tissue compartments from both multishell [3] and single shell [4] diffusion images. We demonstrate that by combining these modern methods with MRI simulation based on the use of the Bloch equations [5], synthetic images can be produced that are of sufficient quality to be used in place of genuine structural images in some standard diffusion tractography analyses. In many circumstances, this may eliminate the need to re-acquire motion-corrupted structural images or to reject participants from analysis due to poor quality structural images.

## 3 Methods

We first describe our proposed method to simulate structural MRIs using diffusion data, then describe *in-vivo* experiments designed to assess this method.

### 3.1 Proposed Method

Our proposed method, summarised in Figure 1, requires that multi-tissue fibre orientation dispersion (FOD) maps have been calculated from diffusion MR data. These maps can be calculated from both single-shell and multi-shell diffusion acquisitions using standard tools without reliance on structural MR images. An example of this process is described in Section 3.2.2.

**Figure 1.**
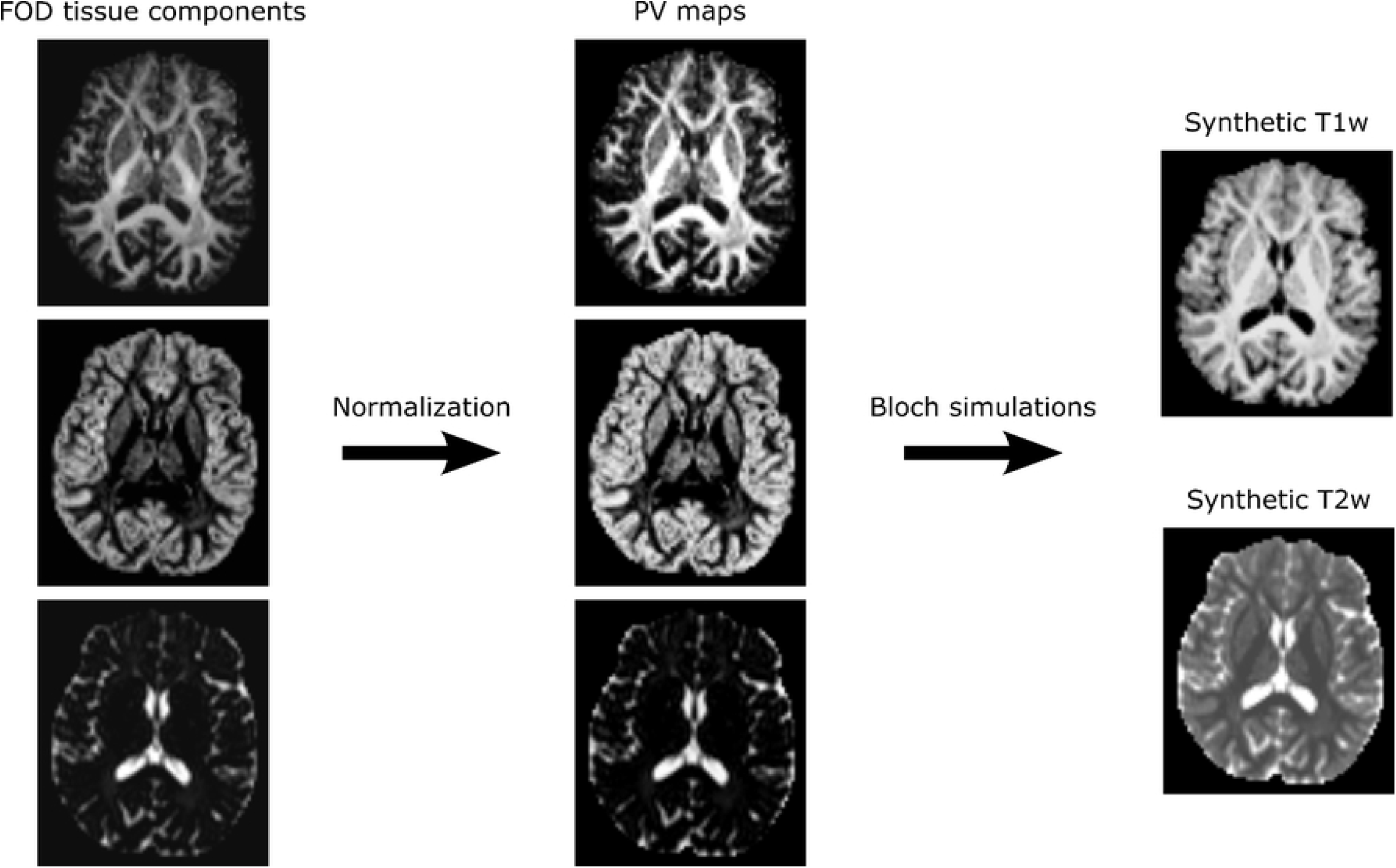
Summary of the proposed MRI synthetization method. After extracting the tissue components of the multi-tissue fibre orientation maps (FOD), partial volume (PV) maps are computed and used to generate synthetic T1w and T2w contrast with Bloch equations-based simulations.

Once calculated, the white matter (WM), grey matter (GM) and cerebrospinal fluid (CSF) tissue components of the FOD maps are extracted and used to generate partial volume (PV) maps as follows:

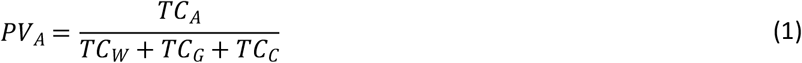

where *PV_A_* and *TC_A_* indicate the partial volume and the FOD tissue component of tissue *A*, respectively, and *TC_W_, TC_G_* and *TC_c_* indicate the WM, GM and CSF tissue components of the FOD. The analytical solution of the Bloch equations for a selected sequence can then be applied to these PV maps to synthesize structural MR images. In the present study, we focus on T1-w and T2-w MRI signals which can be simulated using the following equations:

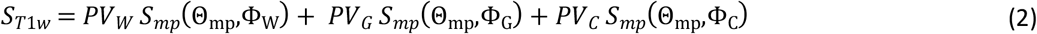

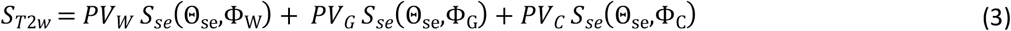

where *S_mp_* and *S_se_* are the analytical solutions of the Bloch equations for the MPRAGE [6] and Spin Echo [7] sequences; Θ_*mp*_ and Θ_*se*_ represent the sequence parameters of the MPRAGE and spin echo sequences; and Φ_*W*_,Φ_*G*_, and Φ_*C*_ correspond to the magnetic properties of the WM, GM and CSF tissues, respectively.

Utilisation of equations 2 and 3 requires selection of scanning parameters and tissue magnetic properties. In the current study, the T1w scans were simulated with the ADNI MPRAGE sequence parameters (*α* = 9°, *TE* = 2.9 *ms, TI* = 900 *ms, TR* = 2300 *ms*) [8]. Similarly, the T2w spin-echo scans were simulated with standard sequence parameters used in clinical imaging (*TE* = 80 *ms, TR* = 4500 *ms*) [9]. The tissue magnetic properties used to simulate the T1w and T2w signals were measured in previous studies conducted at 3T and are summarized in Table 1 [10–12].

**Table 1.** *Magnetic properties of white matter (WM), gray matter (GM) and cerebrospinal fluid (CSF) used for the T1w and T2w signal simulations. These magnetic properties were measured in previous studies conducted at 3T* [10–12].

|  | WM | GM | CSF |
| --- | --- | --- | --- |
| $T1$ (ms) | 830 | 1330 | 4000 |
| $T2$ (ms) | 80 | 110 | 1000 |
| $T2^*$ (ms) | 53 | 66 | 250 |
| $\rho^1$ | 0.7 | 0.8 | 1.0 |

<sup>1</sup>The reported proton densities ( $\rho$ ) are relative measures compared to the CSF proton density.

### 3.2 *In-vivo* experiments

We applied the proposed method to several diffusion datasets in order to assess the quality of the resulting images in terms of qualitative appearance, quantitative tissue contrast, and fitness for use as an integral component of a diffusion tractography pipeline.

#### 3.2.1 Input Data

Three datasets, named herein as the ‘Human Connectome Project Multishell’ (HCP-M), ‘Human Connectome Project Single Shell’ (HCP-S) and ‘Hospital’ datasets, were used in the current study. Scans in these datasets were mostly a convenient sample which had been preprocessed as part of a recent tractography study [13].

The HCP-M dataset included 10 healthy participants from the Human Connectome Project Young Adults 1200 Release [14]. Participants had been scanned twice on a Siemens Skyra Connectom scanner (Siemens Healthcare, Erlangen, Germany) with a 32-channel head coil. We used the preprocessed [15] T1w MPRAGE (1 *mm* isotropic; *α*. = 8°; *TE* = 2.14 *ms*; *TI* = 1000 *ms*; *TR* = 2400 *ms*; acquisition time 7m 40s), T2w SPACE (1mm isotropic; *TE* = 565 *ms*; *TR* = 3200 *ms*; acquisition time 8m 24s) data, and preprocessed multishell diffusion data derived from a high quality acquisition (18 @ b=0s/mm^2^; 90 directions @ b=1000s/mm^2^; 90 directions @ b=2000s/mm^2^; 90 directions @ b=3000s/mm^2^; 1.25mm isotropic). For test-retest experiments (See Section 3.2.4), we utilised structural scans from two time points. For other experiments, only the first time point was used. Ethical consent was granted for use of HCP data.

The HCP-S dataset included the same participants as the HCP-M dataset, excepting that raw data were used (see below) for the diffusion data, which were downsampled and volumes were removed such that only a single shell remained (2 @ b=0s/mm^2^; 60 directions @ b=3000s/mm^2^; no directional repeats; 2mm isotropic). This dataset was designed to represent a modest single-shell HARDI acquisition often seen in the literature.

The Hospital dataset consisted of 12 neurosurgical patients scanned once on a Siemens Prisma scanner with a 64-channel head coil at the Herston Imaging Research Facility in Brisbane, Australia. Images were acquired as part of a clinical trial involving temporal lobe resection for epilepsy (n=2 before surgery, n=6 after surgery), glioma resection (n=2 before surgery), or removal of arteriovenous malformations (n=1 before surgery). This study included acquisition of a T1w MPRAGE scan (1mm isotropic, *α* = 9°; *TE* = 2.9 *ms*; *TI* = 900 *ms*; *TR* = 1900 *ms*; acquisition time 4m 18s) and a modern multishell diffusion acquisition (12 @ b=0s/mm^2^; 20 directions @ b=2000s/mm^2^; 30 directions @ b=1000s/mm^2^; 60 directions @ b=3000s/mm^2^; 2mm isotropic; acquisition time 11m 30s) that is in use in a variety of studies [16–18]. Written informed consent was provided by all patients and the study approved by the Royal Brisbane and Women’s Hospital (RBWH) Human Research Ethics Committee.

#### 3.2.2 Diffusion MRI Processing

Diffusion data from the HCP-S and Hospital datasets were processed automatically using the CONSULT neurosurgical planning pipeline [13]. This pipeline prepared raw diffusion data for tractography using standard libraries and algorithms without requiring associated structural images. Steps included denoising via MRtrix 3.0 [19], removal of motion-corrupted volumes, brain mask calculation using MRtrix’s dwi2mask, and eddy-current distortion correction using a combination of FSL’s topup (http://fsl.fmrib.ox.ac.uk/fsl/fslwiki/TOPUP) and eddy_cuda 8.0. Subsequently, intensity inhomogeneities were corrected using N4-ITK [20]. A final brainmask was then recalculated using MRtrix’s dwi2mask in conjunction with simple morphological operations.

Diffusion data from the HCP-M dataset were downloaded from the HCP server in their minimally preprocessed form. This preprocessing included correction for b0 intensity inhomogeneities, EPI distortion, eddy currents, head motion, gradient non-linearities, as well as reorientation and resampling to 1.25 mm isotropic. Brainmasks for these data were calculated in the same way as for HCP-S and Hospital datasets.

Tissue response functions were calculated using an unsupervised method [21] and FOD maps were calculated for white matter, grey matter, and cerebrospinal fluid using either multishell multitissue contrained spherical deconvolution (for multishell images) [3] or Single-Shell 3-Tissue constrained spherical deconvolution [4] (https://3Tissue.github.io; for HCP-S images). Fractional anisotropy (FA) maps were also calculated from the preprocessed data using MRtrix.

Synthetic T1w and T2w scans were generated from the resulting FOD maps, for all the three datasets, using the method described in Section 3.1.

#### 3.2.3 Contrast measurements

One standard means of assessing structural image quality is measuring intensity contrasts between brain tissues. Here, the brain-tissue contrasts of the *in-vivo* and synthetic structural images were measured using the following equation:

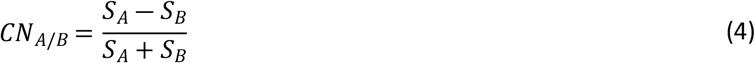

where *S_A_* and *S_B_* refer to the mean signal intensity of tissues *A* and *B*, respectively. The contrast between brain tissues was measured in regions of interest (ROI) manually drawn in the corpus callosum for WM, caudate nucleus for GM and lateral ventricles for CSF. Each ROI comprised at least 20 contiguous voxels. To ensure that the contrast was comparable between scans with different resolutions, specific care was taken to avoid the inclusion of partial volume voxels within the ROIs.

#### 3.2.4 Comparative Performance in a Diffusion Tractography Pipeline

We tested whether the synthetic images were of sufficiently high quality to be used in a standard Freesurfer-based diffusion tractography pipeline, described below. Similar to other tractography-generating processes, this pipeline required structural images for parcellation and diffusion images for tractography. For each subject, we ran this pipeline using (A) *in-vivo* structural scans acquired during the same scan session as the diffusion data (all datasets); (B) *in-vivo* structural scans acquired during a different scan session as the diffusion data (‘*in-vivo* repeat’; HCP-M and HCP-S only); or (C) the synthetically generated structural scans (all datasets). The diffusion FOD image did not differ between each subject’s acquisitions. To determine the influence of the structural scan on reproducibility of this pipeline, we compared these subjects’ acquisitions in terms of both (1) spatial overlap of binarised tractography (Dice score) and (2) differences in FA sampled from this tractography. The former measure was to ascertain how synthetic structural images could impact the tractography itself. The latter was conducted to assess the practical significance of any error introduced by synthetic structural images, as the sampling of FA is a common use-case for tractography.

All image processing was fully automated to avoid biasing results; no manual correction or process re-running was performed for any processing stage. Parcellation of structural images was performed using Freesurfer 6.0 [22]., Both the T1w and T2w images belonging to the HCP-M and HCP-S datasets were provided to Freesurfer. However, only the T1w images were utilised for the Hospital dataset because *in-vivo* T2w scans were not available.

When structural images were acquired *in-vivo*, the resulting parcellation was spatially normalized to the diffusion space by affinely registering the T1w scan to either the FA image (HCP-M and HCP-S) or mean ‘b0’ image (Hospital dataset) using ANTS 2.1 [23]. The mean ‘b0’ image was used for the Hospital dataset as it was already known to produce registrations of better quality for this particular dataset. When structural images were synthetic, no spatial normalization was required because image synthesis naturally creates images in native diffusion space.

The left superior thalamocortical tract was delineated using probabilistic tractography, using labels extracted from the Freesurfer parcellations. Specifically, the thalamus was used as the seeding ROI, while the left primary sensory cortex and left primary motor cortex were combined into a single inclusion ROI. Maximum streamline length was 80 *mm*. Streamlines were acquired using iFOD2 [24] until Tractography Bootstrapping stability criteria [25] were met to ensure the tractogram’s reproducibility was not negatively affected by a low streamline count (minimum streamline count: 10000; min Dice: 0.975; reliability: 0.95; 1.25 *mm* isotropic;*t_bin_*: 0.001 × *n streamlines*). Other parameters were left at default values.

Subject-wise results were compared across runs. To assess overlap between tractograms, each tractogram was converted into a streamline density image at native diffusion resolution, thresholded at the Tractogram Bootstrapping stability threshold (*t_bin_*), and binarised, in line with recommendations for when Tractogram Bootstrapping has been used [25]. The Dice scores between the binarised tractograms of each run were then calculated. To assess reproducibility of microstructural metrics, FA was sampled from each tractogram using MRtrix’s ‘tcksample’ [e.g. 26–28].

## 4 Results

### 4.1 Image Quality and Contrast

An example of PV maps generated from FOD tissue components is presented in Figure 1. Examples of synthetic T1w and T2w scans are presented in Figure 2. Qualitatively, all synthetic scans displayed a similar appearance to their corresponding *in vivo* scan, albeit at the resolution of their diffusion scan (1.25 or 2mm isotropic). Despite this limitation, the borders of individual gyri and subcortical structures were still easily identifiable in all images. Tissue contrast was qualitatively and quantitively similar between *in vivo* and synthetic images, despite small differences in acquisition and simulation protocols (Tables Table 2 and Table 3). Specifically, the synthetic T1w scans were characterized by good suppression of the CSF signal and by a high contrast between grey matter, white matter, and CSF, as is typical for MPRAGE acquisitions. Similarly, the synthetic T2w scans displayed comparable tissue contrasts to those of the *in-vivo* T2w scans, including the typical hyper-intensity of the CSF signal. Synthetic images derived from single and multi-shell diffusion scans produced similar tissue contrast, despite relying on meaningfully different diffusion FOD algorithms and resolutions. Notably, images were also realistic for patients with tumours and post-surgical cavities.

**Figure 2.**
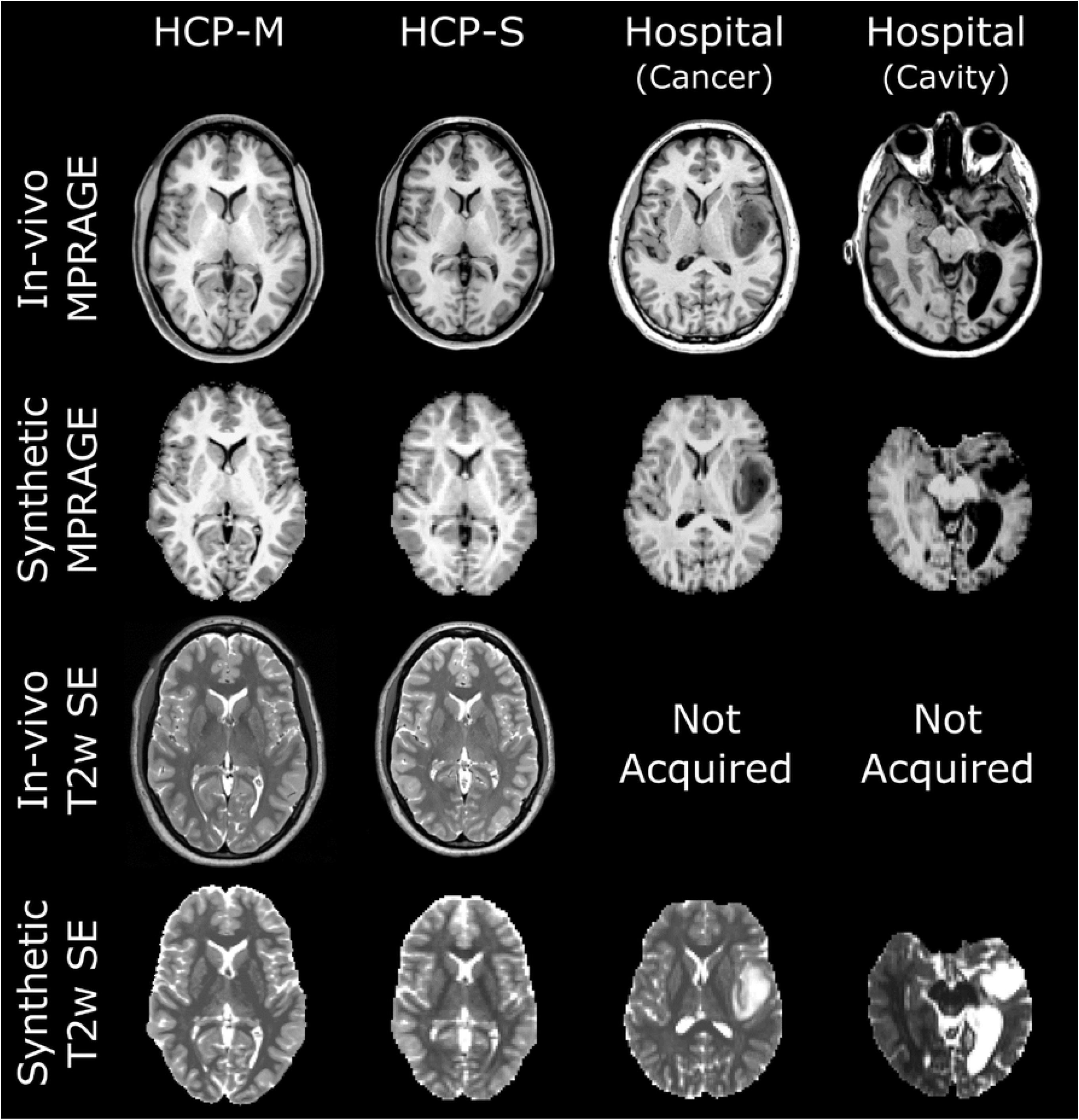
In-vivo and synthetic T1w and T2w images obtained from the HCP-M (far left), HCP-S (middle left) and Hospital (middle and far right) datasets. The synthetic T1w images display contrasts qualitatively similar to that of the in-vivo scans for all datasets, even in presence of tumours (middle right) and post-surgical cavities (far right). The T2w contrast was similar between the synthetic and in-vivo scans for the HCP-M and HCP datasets, including the expected hyper-intensity of the cerebrospinal fluid signal. For the Hospital dataset, the in-vivo T2w images were not acquired but synthetic images provided similar contrast to that which could be expected from an in-vivo scan, including in the areas of pathology.

**Table 2.** Mean brain tissue contrast measured for in-vivo and synthetic T1w scans on the Hospital, HCP-M and HCP-S datasets. HCP datasets used identical in-vivo structural images and so appear together in one row. The minimum and maximum contrast values are shown in parenthesis. The contrasts measured for the synthetic scans are similar to the contrasts measured for the in-vivo scans, despite slight differences between the imaging protocols.

|  | WM/GM | WM/CSF | GM/CSF |
| --- | --- | --- | --- |
| *In-vivo* Hospital | 0.14 (0.12-0.15) | 0.59 (0.56-0.65) | 0.49 (0.46-0.55) |
| *In-vivo* HCP | 0.12 (0.09-0.15) | 0.56 (0.54-0.59) | 0.47 (0.45-0.49) |
| Synthetic Hospital | 0.09 (0.07-0.12) | 0.56 (0.52-0.60) | 0.49 (0.47-0.52) |
| Synthetic HCP-M | 0.10 (0.07-0.12) | 0.50 (0.39-0.60) | 0.43 (0.31-0.55) |
| Synthetic HCP-S | 0.13 (0.10-0.15) | 0.50 (0.42-0.58) | 0.40 (0.33-0.48) |

**Table 3.** Mean brain tissue contrast measured for in-vivo and synthetic T2w scans on the Hospital, HCP-M and HCP-S datasets. The minimum and maximum contrast values are shown in parenthesis. The contrasts measured for the synthetic scans are similar to the contrasts measured for the in-vivo scans, despite the use of different magnetic resonance sequences to acquire and simulate the signals.

|  | WM/GM | WM/CSF | GM/CSF |
| --- | --- | --- | --- |
| *In-vivo* HCP | 0.16 (0.12-0.21) | 0.53 (0.50-0.57) | 0.41 (0.37-0.44) |
| Synthetic Hospital | 0.15 (0.11-0.19) | 0.58 (0.56-0.59) | 0.47 (0.44-0.48) |
| Synthetic HCP-M | 0.16 (0.13-0.18) | 0.58 (0.55-0.59) | 0.46 (0.42-0.49) |
| Synthetic HCP-S | 0.19 (0.16-0.23) | 0.55 (0.51-0.58) | 0.40 (0.36-0.44) |

### 4.2 Comparative Performance in a Diffusion Tractography Pipeline

Freesurfer succeeded for all but one participant. This participant, from the Hospital dataset, presented with a glioma in the left temporal lobe, preventing adequate Freesurfer performance when provided with either *in-vivo* or synthetic input data (Figure 2, middle-left column). A qualitative assessment indicated that Freesurfer parcellations obtained from synthetic datasets were similar to those obtained from *in-vivo* data, as shown in Figure 3. Notably, near the white matter/grey matter interface, from which tractography is typically seeded, Freesurfer parcellations were qualitatively similar between corresponding synthetic and *in vivo* images. However, the synthetic images at 2mm resolution often produced inaccurate segmentation of the pial surface by Freesurfer (Figure 3, Table S1).

**Figure 3.**
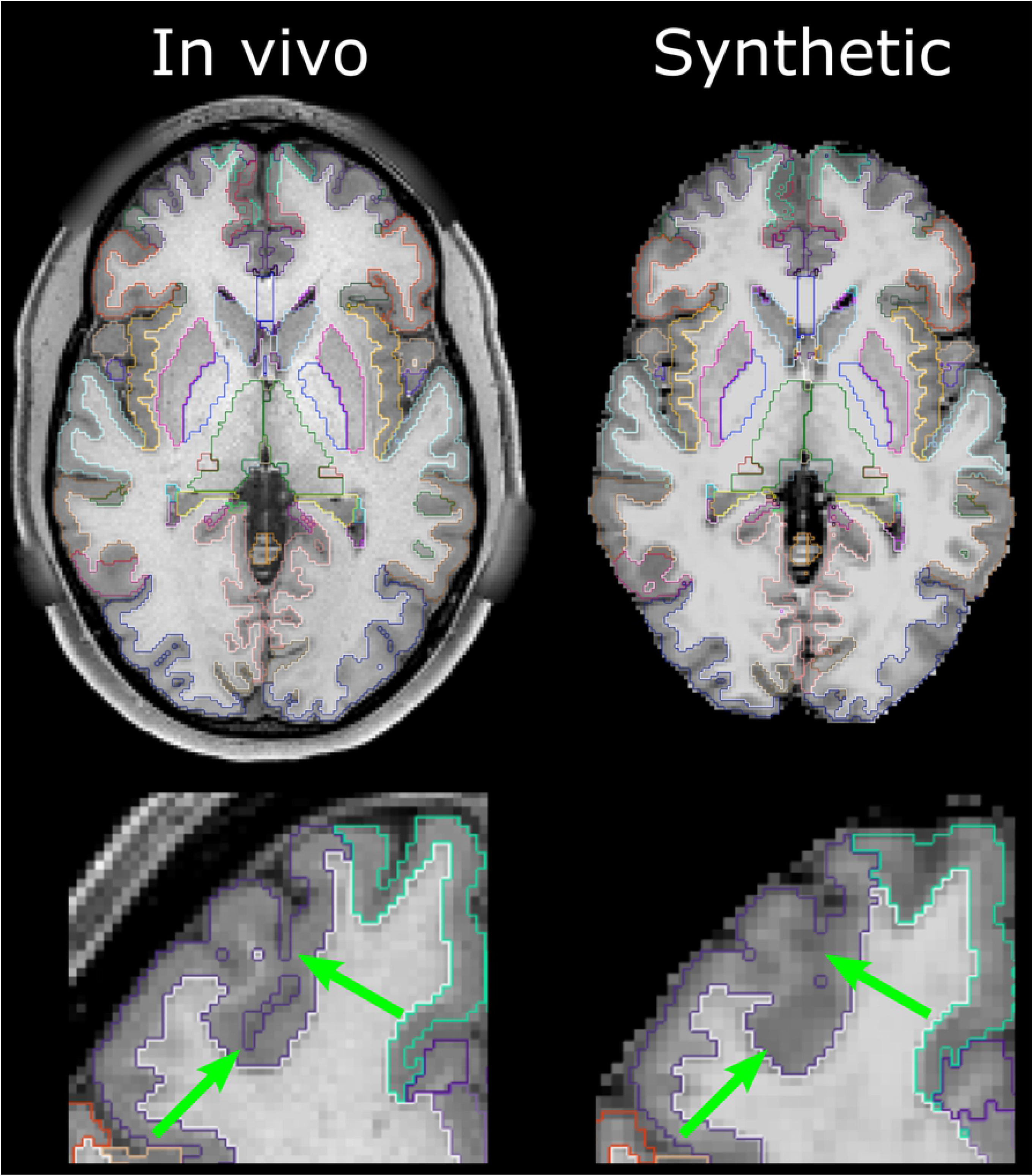
Example of Freesurfer parcellations obtained from the in-vivo (left) and synthetic (right) scans of the Hospital dataset. The bottom row shows a magnified section of the frontal lobe. A qualitative assessment indicated that the synthetic parcellations were similar to the in-vivo parcellations in subcortical structures and near the grey-matter/white-matter interface from which tractography is typically seeded. However, the lower resolution provided by the synthetic data (2 mm isotropic) reduced the accuracy of the pial surface segmentation, as highlighted by the green arrows.

Tractography was successfully performed for all the datasets for which Freesurfer parcellation did not fail. Tractography was qualitatively similar between in*-vivo*- and synthetic-structural pipelines (Figure 4). Median Dice coefficients for binarised tractography between the *in-vivo*- and synthetic-structural pipelines were 0.93 (HCP-M), 0.93 (HCP-S), and 0.90 (Hospital) with respective worst-case performances of 0.90, 0.90, and 0.88 (Figure 5). Structural scans were available from a second time point for all HCP datasets. The tractography pipeline was repeated using the same diffusion data but with these alternative *in-vivo* structural scans. These *in-vivo* vs *in-vivo*-repeat comparisons demonstrated similar median Dice coefficients to, and poorer worst-case performance than, the aforementioned *in-vivo* vs *synthetic* comparisons (Bonferroni-corrected Wilcoxon Signed-Rank Tests, both p>0.1; See Figure 5). This poorer worst case performance may be explained by variations in the quality of registration between structural and diffusion scans: a step not required by the synthetic pipeline which guarantees perfect alignment between modalities.

**Figure 4.**
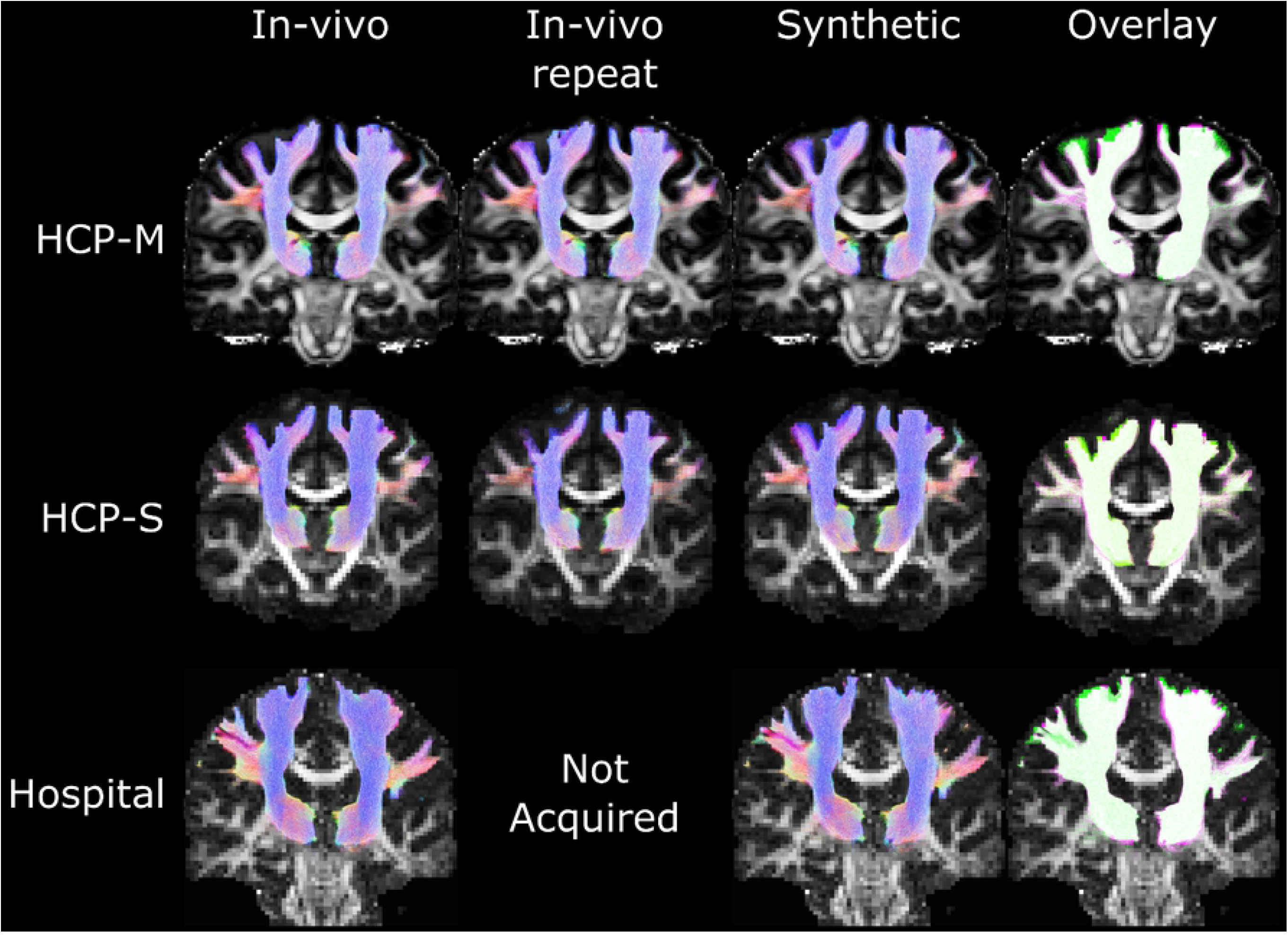
Example tractography from the in-vivo and synthetic pipelines. The rightmost column shows overlap of the in-vivo tractography (from the leftmost column; green) with the synthetic tractography (fuchsia; white indicating overlap). Tractography densities showed clear correspondences between in vivo and synthetic pipelines for HCP-S, Hospital, and HCP-M datasets. The datasets shown here were selected at random.

**Figure 5.**
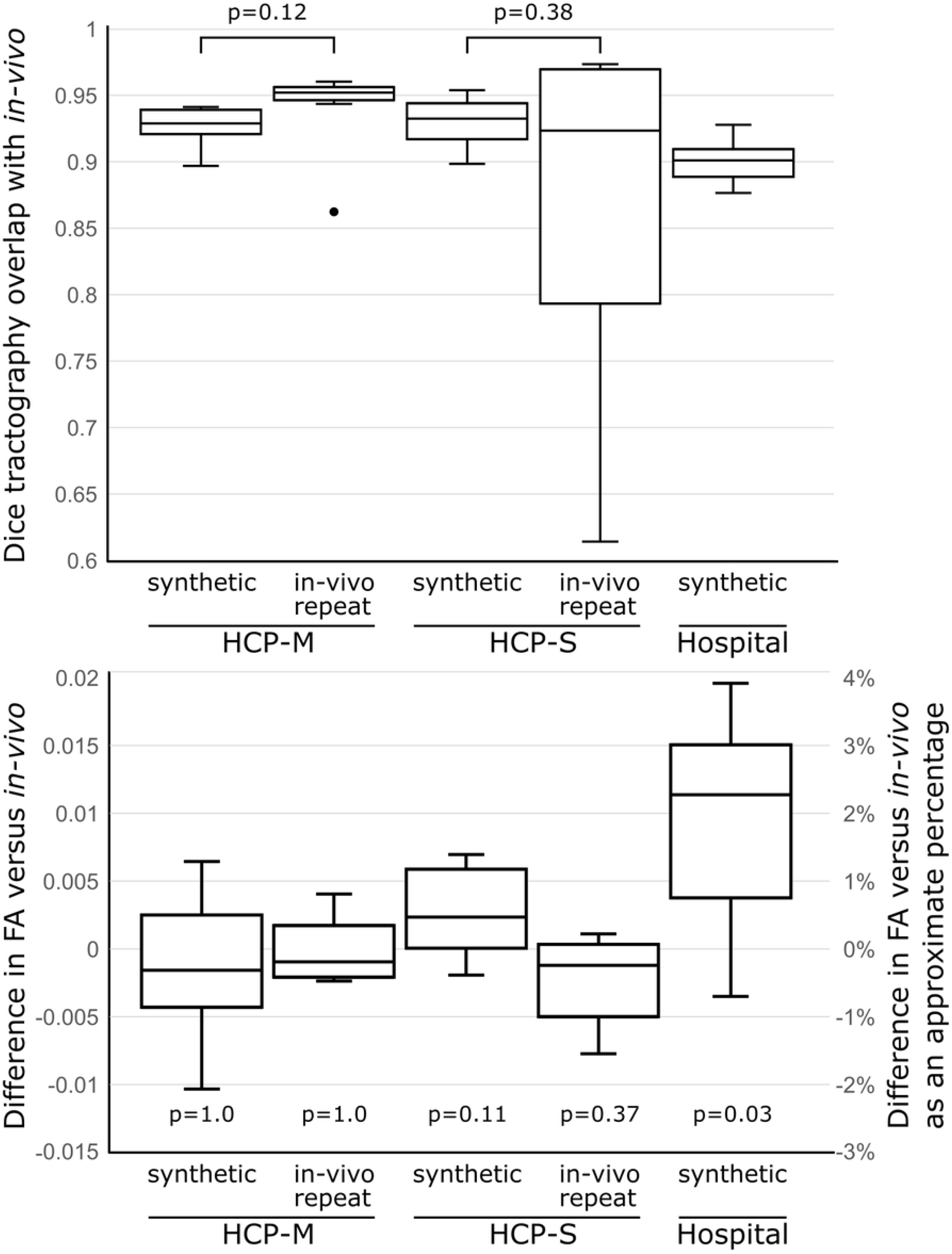
The tractography pipeline was run first using genuine (‘in-vivo’) diffusion and structural MR images, and then run again using the same diffusion data but with either a synthetic structural image (‘synthetic’) or a structural scan acquired during another scanning session (‘in-vivo repeat’). Top: Dice scores for overlap of tractography between runs using in-vivo data and either synthetic or in-vivo repeat data. Median dice scores did not differ significantly between synthetic and in-vivo repeat runs after correction for multiple comparisons. Bottom: Mean tract FA values sampled from the tractography, compared with the in-vivo tractography. Differences between the in-vivo versus synthetic or in-vivo-repeat runs were below the level of practical significance. Approximate percentages were calculated by dividing differences by the mean FA of all datasets (0.52). Displayed p-values are corrected for multiple comparisons.

FA values sampled from tractography differed significantly between the synthetic-structural pipeline and *in-vivo*-structural pipeline only for the Hospital dataset (Bonferroni-corrected Wilcoxon Signed-Rank tests versus the *in-vivo*-structural pipeline FA values: HCP-M Synthetic, p = 1; HCP-M *in-vivo* repeat, p = 1; HCP-S Synthetic, p = 0.11; HCP-S *in-vivo* repeat 0.37; Hospital Synthetic, p = 0.02). Although statistically significant, this median difference was only 0.01, or 2.2%, which is well below both inter-group and longitudinal differences typically reported in imaging studies [26–28].

## 5 Discussion

Motion artefacts are commonplace in MRI acquisition, particularly with children and cognitively impaired persons [2,29]. Diffusion imaging can be easily ‘scrubbed’ of motion affected volumes, typically leaving usable data.However, the same is not true for structural images which are critical to most diffusion analyses. As such, motion in structural images can lead to disproportionate data loss [2,29]. To alleviate this issue, we have proposed a means by which synthetic structural images can be generated from diffusion MR images. We are aware of one previous report [30] in which a single diffusion-MR derived segmentation was normalised voxel-wise and multiplied by experimentally derived values in order to generate an image qualitatively similar to a T1w image. We have expanded on this idea by using a well-established physical model that allows precise scanning parameters to be simulated; this is demonstrated here by the synthesis of specific T1w MPRAGE and T2w spin-echo sequences from both single-shell (HCP-S dataset) and multi-shell (HCP-M dataset, Hospital dataset) diffusion scans. We demonstrated that the synthetic images generated for these 32 participants, including those displaying pathology, were both visually convincing and had tissue contrasts in line with images acquired *in-vivo*.

To demonstrate the utility of this method, we performed Freesurfer-driven tractography for 32 participants. Freesurfer is a well-established program and is regularly relied on for diffusion analyses but can, like many packages, fail if low-quality images are provided. Here, Freesurfer was able to segment brains successfully in all instances for both *in-vivo* and synthetic data excepting for one subject presenting with a glioma for which it failed with both real and synthetic images. Due to the lower resolution of the synthetic data, the pial surface was not always as well defined in the synthetic-image parcellation as in the *in-vivo* image parcellation. However, this is unlikely to be of serious concern for tractography-based studies which are more reliant on an accurate grey-matter/white-matter interface, which was generally of high quality here (Figure 3). Indeed, tractography derived from *in-vivo* structural and synthetic-structural parcellations overlapped well (median Dice ≥ 0.90) for all three types of diffusion acquisition tested in this study. A common use for tractography is to identify a tract from which microstructural measures, such as FA, can be sampled. Here, FA sampled from tractography of the *in-vivo* and synthetic-image pipelines differed well below the level of practical significance for most forms of analysis (Figure 5).

An alternative to the proposed method, available only to longitudinal diffusion MRI studies, is to replace a corrupted structural image with another acquired at another time point. We tested the efficacy of this alternative for the HCP-S and HCP-M datasets by running the Freesurfer-based pipeline with T1w and T2w scans acquired at a second time point. In general, this did not produce meaningfully better results, in terms of either Dice scores or FA, than using a synthetic image. Visual inspection suggested that performance of this *in-vivo-repeat* run sometimes suffered from slight mis-registrations between the real structural images and diffusion images, reminding that the negative impact of the lower-resolution synthetic images is somewhat counter balanced by their guaranteed perfect alignment to the diffusion image.

One strength to our method is a reliance on a well-established physical model that makes a relatively small number of assumptions. While hand-tuned or machine-learning based approaches could in principle generate similar outputs, these are inherently tuned to their training data, which sometimes leaves general use uncertain – for example, when presented with lesions, low quality data, or high resolution inputs. By contrast, our proposed method is resolution independent and, as demonstrated here, was able to generate convincing images from data acquired on two scanners, using two different tissue response function methods, and three different acquisition parameters. It also was able to produce convincing and usable images where pathology and post-surgical cavities were present without requiring tuning of any kind.

Importantly, the proposed method is intended to enable diffusion MRI analyses of white matter; it is not intended to produce images for clinical interpretation nor for other types of analyses that would normally rely on precise tissue-contrast for certain pathologies or sharp pial surfaces. For this intended use-case, the major limitation of the proposed method is that the synthetic images produced are at the resolution of the diffusion data. Although it is possible to apply up-sampling and super-resolution techniques, we have not attempted to do so here because the synthetic images were of sufficient quality to obtain adequate quality parcellations and diffusion metrics. It is also worth considering that the proposed method provides some insurance against motion-corrupted structural images, therefore, scan time does not necessarily have to be pre-allocated for potential re-acquisition of motion-affected structural scans. This, in turn, may provide some studies with additional scan time that can potentially be used to modestly improve the resolution of their diffusion data.

In conclusion, we have presented a simple and physically constrained means of synthesising structural images with customisable acquisition parameters from diffusion MRI. These images are of adequate quality to be used with standard parcellation tools, allowing analysis of diffusion data that would otherwise be impossible due to motion-corrupted or non-acquired scans.

## 6 Data reporting

The code that supports the findings of this study is openly available in the FLAWS-Tools repository at https://github.com/jerbeaumont/FLAWS-Tools.git (https://doi.org/10.5281/zenodo.3993247). Please note that the code will be released in this repository once this article will be accepted for publication.

The sharing of patients’ data from the Hospital dataset was not approved by the Royal Brisbane and Women’s Hospital (RBWH) Human Research Ethics Committee. Therefore, the images from the Hospital dataset cannot be shared. For the HCP dataset, the FOD and T1 images (both real and synthetic) will be made available upon paper acceptance. All numerical results reported in this paper are detailed per subjects in Annexes S1 and S2.

## 7 Acknowledgements

This research was partially supported by an Advance Queensland Research Fellowship (R-09964-01) and by the ‘Region Bretagne’ (France). Data were provided in part by the Human Connectome Project, WU-Minn Consortium (Principal Investigators: David Van Essen and Kamil Ugurbil; 1U54MH091657) funded by the 16 NIH Institutes and Centers that support the NIH Blueprint for Neuroscience Research; and by the McDonnell Center for Systems Neuroscience at Washington University. The authors are grateful to the Dr Katie McMahon, Ms Clare Berry, and the team at the Herston Imaging Research Facility for their generous scan time allowances, and technical support acquiring some of the data presented here. The authors acknowledge the facilities of the National Imaging Facility, a National Collaborative Research Infrastructure Strategy (NCRIS) capability, at the Herston Imaging Research Facility.

